# The Dynamic Landscape of Transcription Initiation in Yeast Mitochondria

**DOI:** 10.1101/2019.12.16.877878

**Authors:** Byeong-Kwon Sohn, Urmimala Basu, Seung-Won Lee, Hayoon Cho, Jiayu Shen, Aishwarya Deshpande, Smita S. Patel, Hajin Kim

## Abstract

Controlling efficiency and fidelity in the early stage of mitochondrial DNA transcription is crucial for regulating cellular energy metabolism. Studies of bacteriophage and bacterial systems have revealed that transcription occurs through a series of conformational transitions during the initiation and elongation stages; however, how the conformational dynamics progress throughout these stages remains unknown. Here, we used single-molecule fluorescence resonance energy transfer techniques to examine the conformational dynamics of the two-component transcription system of yeast mitochondria with single-base resolution. We show that, unlike its single-component homologue in bacteriophages, the yeast mitochondrial transcription initiation complex dynamically transitions between closed, open, and scrunched conformations throughout the initiation stage, and then makes a sharp irreversible transition to an unbent conformation by promoter release at position +8. Remarkably, stalling the initiation complex revealed unscrunching dynamics without dissociating the RNA transcript, manifesting the existence of backtracking transitions with possible regulatory roles. The dynamic landscape of transcription initiation revealed here suggests a kinetically driven regulation of mitochondrial transcription.

## Introduction

Recent studies of transcription machinery have found that transcription initiation is not a unidirectional process that invariably leads to elongation, but is rather a stochastic process that involves divergent pathways such as abortive initiation, backtracking, and pausing in addition to the progression to elongation^1-6^. These findings suggest that clearing initiation and progressing to elongation might be rate-determining steps that control transcription efficiency, and they have been suggested to be targets of transcription regulation^3,7-9^. Transcription systems across lineages with different levels of complexity show highly variable initiation efficiencies that depend on specific DNA elements^10,11^. The initiation stage and transition to elongation are regulatory targets for many proteins and small molecules^8,12^. Thus, investigating conformational dynamics during the early stage of transcription is crucial to understanding the regulation of transcription efficiency and fidelity.

While multi-subunit RNAPs have been studied in great detail, we lack such deep understanding of the mitochondrial transcription systems that share homology with bacteriophage transcription systems. T7 RNA polymerase (RNAP), the single-subunit transcription machinery of bacteriophages, is homologous to the yeast mitochondrial RNAP (Rpo41) and the human mitochondrial RNAP (POLRMT)^13-15^. Transcription systems in bacteriophages and mitochondria also share common promoter recognition mechanisms^15^. While T7 RNAP does not require additional proteins to initiate transcription, yeast mitochondrial transcription requires the initiation factor Mtf1, which is responsible for stabilizing open promoter regions^6,16-18^. Similarly, human mitochondrial transcription requires the initiation factors TFAM and TFB2M^19-21^. Mtf1 is structurally and functionally homologous to TFB2M^19,22^, and is functionally similar to the bacterial sigma factor^23-25^. To regulate transcription efficiency at various promoters, these transcription systems have developed a set of molecular mechanisms, in which they share several key features. Both bacteriophage and bacterial RNAPs show scrunching of the downstream DNA into the active site as the RNA-DNA hybrid and the transcription bubble grow during initiation. After a stable full-length RNA-DNA hybrid is formed, the RNAP releases upstream promoter contacts and the initiation bubble collapses to drive the transition into elongation^26-32^. A similar mechanism is also thought to exist in higher organisms^33,34^. More recently, branching between competing pathways and pausing during initiation have been observed in both bacteriophages and bacteria and may play crucial roles in regulating transcription activity^1,3,4,35^.

In yeast mitochondria, transcription initiates with the assembly of Rpo41 and Mtf1 at conserved promoter sequences located at positions −8 to +1 relative to the transcription start site. The 2-aminopurine (2AP) fluorescence and protein-DNA crosslinking experiments have shown that Mtf1 facilitates promoter melting at positions −4 to +2 by trapping the non-template strand^24,25,36^. The structures of T7 and human mitochondrial RNAPs show that the promoter DNA is severely bent around the start-site in the open complex^37,38^. Single-molecule studies of the yeast mitochondrial RNAP have shown that the open promoter dynamically switches between bent and unbent conformations^6^. The way in which the yeast mitochondrial transcription initiation complex (TIC) transitions from initiation to elongation has not been observed directly, but its single-subunit relative, T7 TIC, exhibits a concerted motion of DNA scrunching and rotation during initiation, and the promoter unbends upon transition to the elongation complex^30,32^. During transcription initiation by T7 RNAP, the complex adopts a rather static form, represented by a dominant population of conformation at each step. Ensemble-level and single-molecule studies of the T7 transcription system show that transition to the elongation stage occurs through multiple steps, gradually over positions +8 to +12^30,39,40^.

In this study, we used single-molecule fluorescence resonance energy transfer (smFRET) techniques and ensemble biochemical assays to reveal the complex conformational dynamics of the yeast mitochondrial transcription throughout the initiation stage and transition to the elongation stage. As expected, we found that the DNA template progressively bends and scrunches during initiation; however, in contrast to the T7 transcription system, it continues to show transition between closed, open, and scrunched conformations throughout the whole initiation stage. Intriguingly, reversible scrunching and unscrunching transitions in the stalled complexes revealed complex branching kinetics during transcription initiation. The unscrunching transitions were not necessarily accompanied by dissociation of the RNA transcript and appear to be backtracked initiation complexes. At position +8, a sharp conformational transition was observed, marked by the transformation of the upstream DNA into a stable unbent form. The unbending conformational transition at +8 was then followed by a gradual collapse of the initiation bubble to complete the transition into the elongation stage. These findings are in stark contrast to the T7 transcription system, which exhibits more static initial complexes, and synchronous but a gradual unbending and bubble collapse transition over several base positions to the elongation complex. Thus, our results reveal novel conformational dynamics during initiation by the mitochondrial RNAP that could be vital as checkpoints for regulating mitochondrial DNA transcription efficiency and promoter selection.

## Results

### Transcription initiation progresses through dynamic conformational ensembles

In order to probe the conformational dynamics of the yeast mitochondrial TIC using smFRET techniques, we designed a 50-basepair DNA template containing the yeast mitochondrial promoter sequence and fluorescently labeled at specific positions upstream and downstream of the transcription start site (Fig. 1a). The DNA was extended to position −34 and modified with biotin at the 3’ end of the template strand for surface immobilization. Position −16 of the non-template strand was labeled with Cy5 and position +16 of the template strand was labeled with Cy3 (Fig. 1a and 1b), such that the FRET signal between the dyes reported bending and scrunching of the template in real time. The coding sequence was designed to stall transcription at positions +2, +3, +5, and +6 by using a combination of ribonucleotides and 3’-deoxyribonucleotides (DNA template I; Fig. 1b). Another template (DNA template II) was designed to stall transcription at positions +7 and +8.

**Figure 1.**
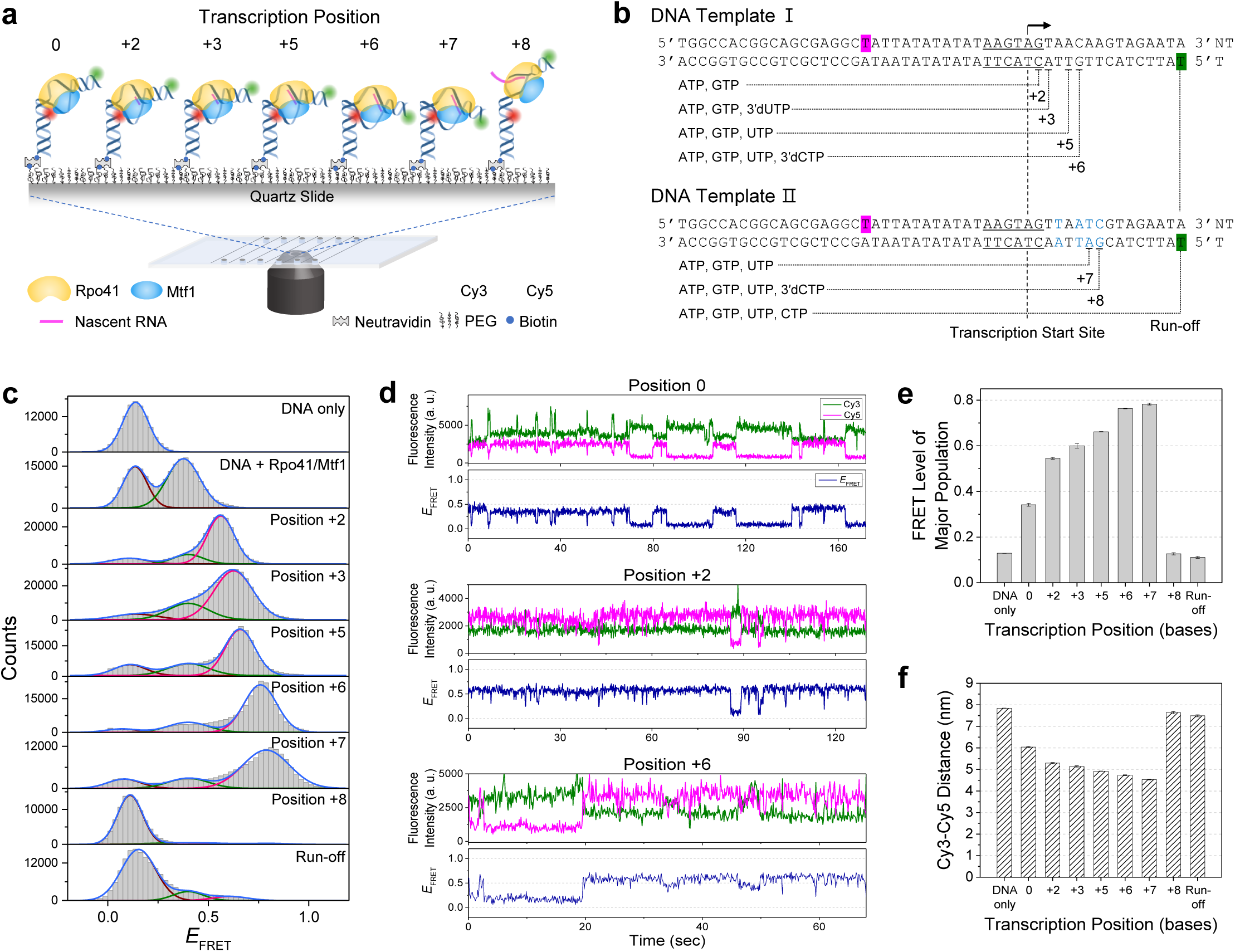
Transcription initiation occurs through dynamic conformational changes of the initiation complex. **a**, Single-molecule measurements of transcription initiation dynamics. The dual-labeled DNA template complexed with Rpo41 and Mtf1 was observed using a total internal reflection fluorescence microscope. **b**, DNA templates used in basepair-wise measurements of initiation complex dynamics. DNA template I could be stalled at positions +2, +3, +5, and +6, while DNA template II, which differed from DNA template I by four basepairs (blue), could be stalled at positions +7 and +8. Both templates were labeled with Cy5 at position −16 of the non-template strand (magenta) and Cy3 at position +16 of template strand (green). The transcription promoter (underscored) and start site (arrow) are indicated. **c**, FRET histograms from single-molecule traces with colocalized Cy3 and Cy5 signals at each stalling position. Histograms were fit to single, double, or triple Gaussian peaks. The brown, green, and magenta curves represent low, mid, and high FRET populations, respectively. **d**, Representative smFRET traces at positions 0, +2, and +6 showing the Cy3 (green) and Cy5 (magenta) signals and the FRET efficiency traces (navy). **e**, The FRET level of the major population in (**c**) shown for each stalling position as the center of the major Gaussian peak. Error bars represent the error in Gaussian fitting. **f**, The Cy3-to-Cy5 distance at each stalling position calculated from the average between major FRET levels from DNA templates I/II (**e**) and I/II NT (Supplementary Fig. 1c). Error bars represent propagation of the errors in FRET levels.

The FRET efficiency distribution at each transcription stalling position was obtained from a large number of single-molecule traces (Fig. 1c). Bare DNA template I exhibited a single peak at low FRET efficiency (*E*_FRET_ = 0.14). When complexed with Rpo41 and Mtf1, the DNA showed two peaks, one at a low *E*_FRET_ (0.14) and the other at a mid *E*_FRET_ (0.38). The low FRET population does not represent bare DNA, because time traces showed dynamic transitions between the low and mid FRET levels (Fig. 1d). Moreover, these transitions were not due to binding and dissociation of proteins, as the same dynamics were observed when unbound proteins were washed out, consistent with a previous report^6^. Such persistent FRET dynamics in DNA complexed with Rpo41 + Mtf1 represent conformational transitions between closed (low FRET) and open (mid FRET) promoter states^6^.

As transcription was progressed to position +2 by equilibrating the TIC of DNA template I with ATP and GTP (0.5 mM each), we observed a higher FRET state (*E*_FRET_ = 0.56) that dominated the trace but exhibited transitions to the previously observed low and mid FRET states (Fig. 1c and 1d). The high FRET state presumably represents the scrunched DNA template at position +2, and our results suggest that the scrunched TIC can become unscrunched, switching to a conformation similar to that of the open promoter state, and can sometimes even switch back to the closed promoter state, presumably following the dissociation of RNA product by abortive initiation. Such scrunching-unscrunching dynamics during transcription initiation were recently observed in bacterial transcription machinery and were suggested to provide a paused checkpoint for abortive and productive RNA synthesis^3^. Upon stalling the TIC at positions +3, +5, +6, and +7, the major FRET population progressively rose to higher FRET levels of 0.63, 0.66, 0.76, and 0.79, respectively. The low and mid FRET populations remained small but switching between the three FRET states persisted (Fig. 1c and 1d). As the concentration of ribonucleotide mix (NTP) was raised at position +7, the high FRET population increased but its FRET level did not change (Supplementary Fig. 1). At a lower NTP concentration of 50 µM, the high FRET population decreased and the TIC exhibited an extended stay at the mid FRET state. At 5 µM NTP, the TIC no longer progressed to the high FRET state. Thus, the high FRET state observed at each stalling position likely represents the major conformation of the TIC stalled at the desired position. Notably, upon stalling the TIC at position +8, the high FRET population disappeared almost completely, and a low FRET population became dominant (Fig. 1c). Under run-off conditions with all ribonucleotides supplied, the FRET distribution was similar to that at position +8.

Overall, these results show that the TIC progresses through a series of dynamic conformational ensembles during transcription initiation. The FRET level of the major population gradually increased upon progression from position +2 to +7, followed by an abrupt drop at position +8 (Fig. 1e). At all steps, the TIC continued to show dynamic transitions between distinct FRET states. Based on the changing FRET distribution, we propose that the TIC gradually scrunches up to position +7, but keeps switching between several conformations. Subsequently, the tension caused by scrunching is suddenly released at position +8, allowing the DNA template to form an extended form (Fig. 1a).

The FRET level does not precisely represent the bending and scrunching motions of duplex DNA, because it is affected by the DNA twisting motion that accompanies the insertion of the downstream DNA into the transcription site. Hence, we generated alternative DNA templates I and II in which the downstream Cy3 label was positioned on the non-template strand instead of the template strand (DNA templates I NT and II NT; Supplementary Fig. 2). Like those generated from DNA templates I and II, the FRET histograms generated from DNA templates I NT and II NT displayed a gradual increase in the FRET level up to position +7 and a sudden drop at position +8 (Supplementary Fig. 2). By taking an average of the FRET levels from these two sets of DNA templates, we obtained the distances (*R*_D-A_) between position −16 of the non-template strand and the center of the downstream DNA at position +16 at each stalling position (Fig. 1f). These distances were compared with those calculated using the crystal structure of the human mitochondrial TIC (PDB: 6ERP; Supplementary Fig. 2)^41^. Because 6ERP provides the structure of the initiation complex at position 0 (IC0), we calculated the distances at other stalling positions by assuming that the angle between the upstream and downstream DNA arms is fixed, and that the downstream DNA only scrunches and twists during initiation. The *R*_D-A_ of IC0 matched well between the smFRET data and the crystal structure but showed discrepancies at further stalling positions; specifically, the crystal structure estimated the distance to be larger (Supplementary Fig. 2). This finding implies that the DNA arms gradually bend during initiation, which places the dye pair in closer proximity. The bending angle at each stalling position calculated from the smFRET data started at 110 degrees, gradually increased to 124 degrees at position +7, and then dropped abruptly to 83 degrees at position +8, presumably representing the less bent form of DNA in elongation complex (EC) (Supplementary Fig. 2).

**Figure 2.**
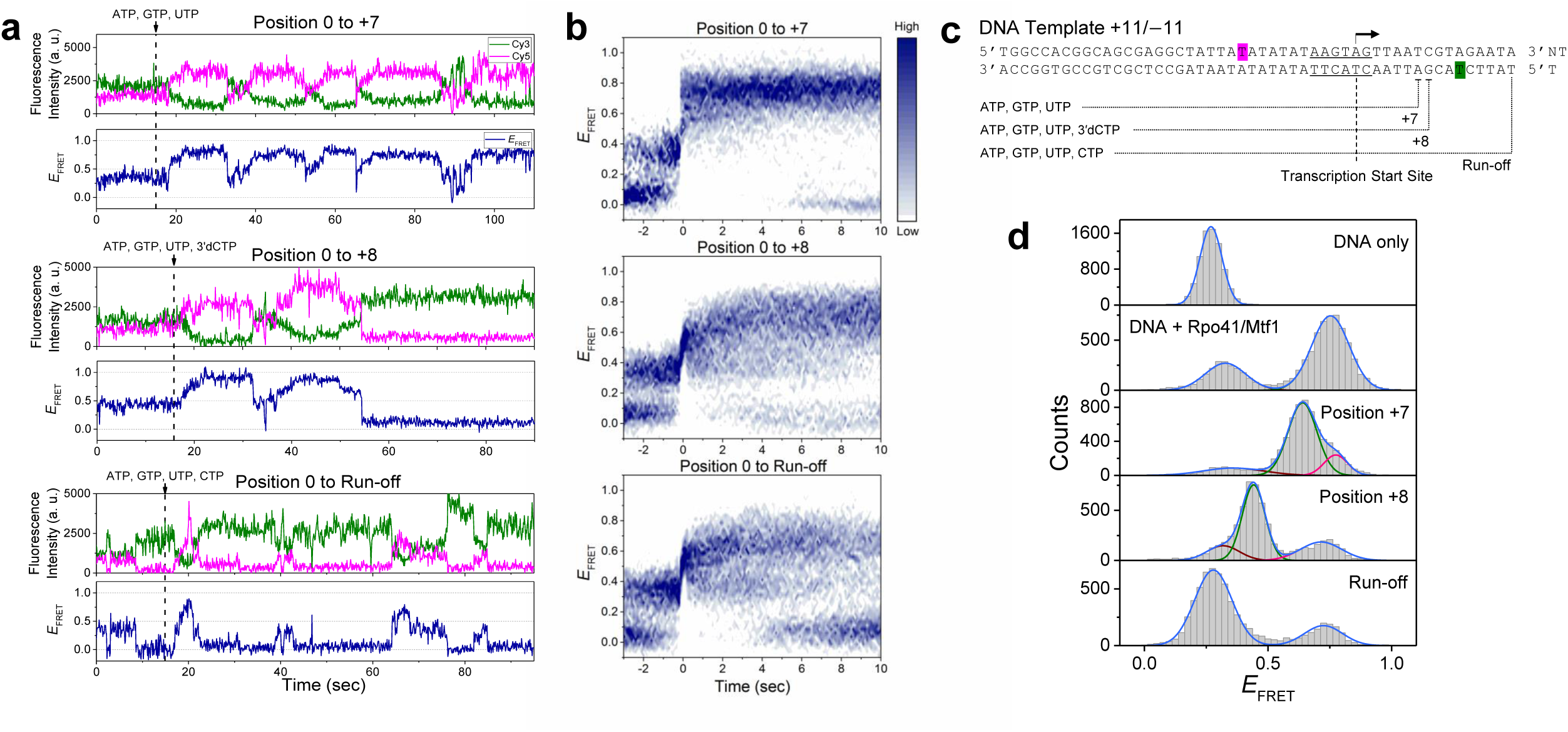
Transition to elongation occurs via a large, abrupt conformational change at position +8. **a**, Representative smFRET traces from flow-in measurements. The point at which combinations of NTPs were added to stall the initiation complex at position +7 or +8 or to enable run-off is shown as a vertical dotted line. **b**, FRET evolution maps constructed by overlaying multiple traces synchronized at the moment the FRET signal reached 0.5 (0 second). 176, 181, and 160 traces were used to generate maps for progression to positions +7 and +8, and run-off, respectively. **c**, Schematic design of DNA template +11/−11 used to distinguish between the conformations of the elongation complex and DNA only. **d**, FRET histograms from DNA template +11/−11. Single, double, or triple Gaussian fitting to each histogram is shown.

### Transition to the elongation stage occurs with large, abrupt conformational changes

To examine the transition from initiation to elongation between +7 and +8, we designed another set of experiments in which we recorded smFRET movies while flowing the ribonucleotide mixture into IC0. Such flow-in measurements allowed us to track the FRET transitions in real time. When the ribonucleotide mixture (0.5 mM each of ATP, GTP, and UTP) was flowed in to progress the TIC to position +7, the initial mid FRET state switched to the high FRET state within several seconds and stayed there for a while. Subsequently, it transitioned back to the low or mid FRET state, followed by another rise to the high FRET state (Fig. 2a). These dynamics continued, presumably reflecting either the scrunching-unscrunching dynamics of the TIC or abortive initiation followed by rounds of re-initiation.

Similar dynamics were observed during progression to position +8 (promoted by flowing in 0.5 mM each of ATP, GTP, UTP, and 3′dCTP), including extended stays at the high FRET state and transitions back to the low/mid FRET state. However, the high FRET state eventually switched abruptly to a prolonged low FRET state (Fig. 2a). The level of this low FRET state matched that of the major population in the equilibrium measurements (Fig. 1c). Therefore, the low FRET state represents the equilibrium structure of EC at position +8 (EC8). Flow-in measurements with all ribonucleotides to promote run-off transcription (0.5 mM each of ATP, GTP, UTP, and CTP) showed a similar pattern of traces, but in this case, the long-lived low FRET state repeatedly returned to the mid and high FRET levels (Fig. 2a).

This finding plausibly represents transcription re-initiation with a new protein complex after run-off synthesis of the previous transcript. The complex stalled at positions +7 and +8 and under run-off conditions showed the same dynamic behaviors at equilibrium as those in the flow-in measurements (Supplementary Fig. 3). However, the FRET dynamics at position +8 were rarely observed due to long stalling at the low FRET state.

**Figure 3.**
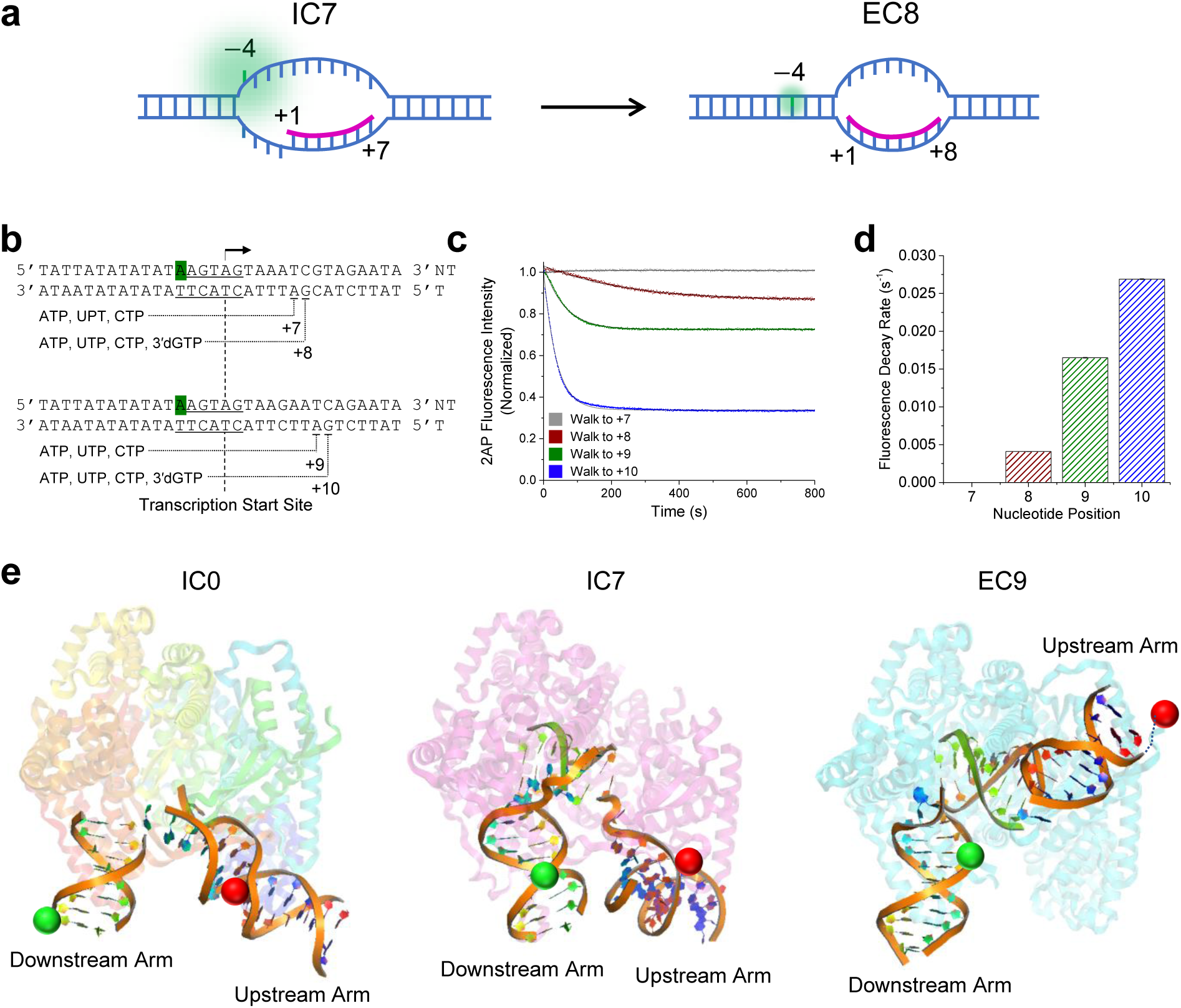
Transcription initiation bubble collapses upon transition to elongation. **a**, Schematic illustration showing IC7 with the initiation bubble and a 7-nt RNA (magenta) annealed to the template DNA from positions +1 to +7. The template bases from −4 to −1 are single-stranded, resulting in a strong fluorescence signal of 2AP at position −4 of the non-template strand (green). At position +8, EC8 is shown where the initiation bubble has collapsed and the −4 to −1 region is reannealed, resulting in the quenching of 2AP fluorescence. **b**, Design of DNA templates used for 2AP fluorescence measurements, to be stalled at positions +7, +8, +9, and +10. **c**, Changes in 2AP fluorescence measured along *in vitro* transcription reactions to the indicated positions, normalized against the initial intensity. The intensity traces were fit to a single exponential decay curve to determine the transition rates. **d**, Fluorescence decay rates measured from (**c**) at different walking positions. **e**, Models of the IC0, IC7, and EC9 structures generated using PyMOL (Schrödinger, USA). IC0 was modeled using PDB 6erp (human mitochondrial RNA polymerase initiation complex), IC7 was modeled using PDB 3e2e (bacteriophage T7 RNA polymerase initiation complex with 7 bp RNA:DNA), and EC9 was modeled using PDB 4boc (human mitochondrial RNA polymerase elongation complex with 9 bp RNA:DNA). The green and red balls represent the Cy3 and Cy5 fluorophores at positions +11 and −11, respectively. The double-stranded DNA and RNA:DNA hybrid (RNA in green) is highlighted as bound to the protein in the background.

Under run-off conditions, the Cy3 signal increased immediately after the abrupt drop in the FRET level (Supplementary Fig. 4). This protein-induced fluorescence enhancement^42^ accompanying run-off RNA synthesis was attributable to Rpo41 + Mtf1 translocating over the Cy3 fluorophore in the downstream DNA. Comparison of the fluorescence intensity histograms at stalling position +2 and under run-off conditions confirmed the protein-induced fluorescence enhancement effect by revealing the presence of an additional peak at a higher intensity for run-off transcription (Supplementary Fig. 4). These results indicate that Rpo41 + Mtf1 translocates all the way to the end of the DNA template when supplied with all ribonucleotides, further supporting that the abrupt drop in the FRET level reflects a transition to the elongation stage.

**Figure 4.**
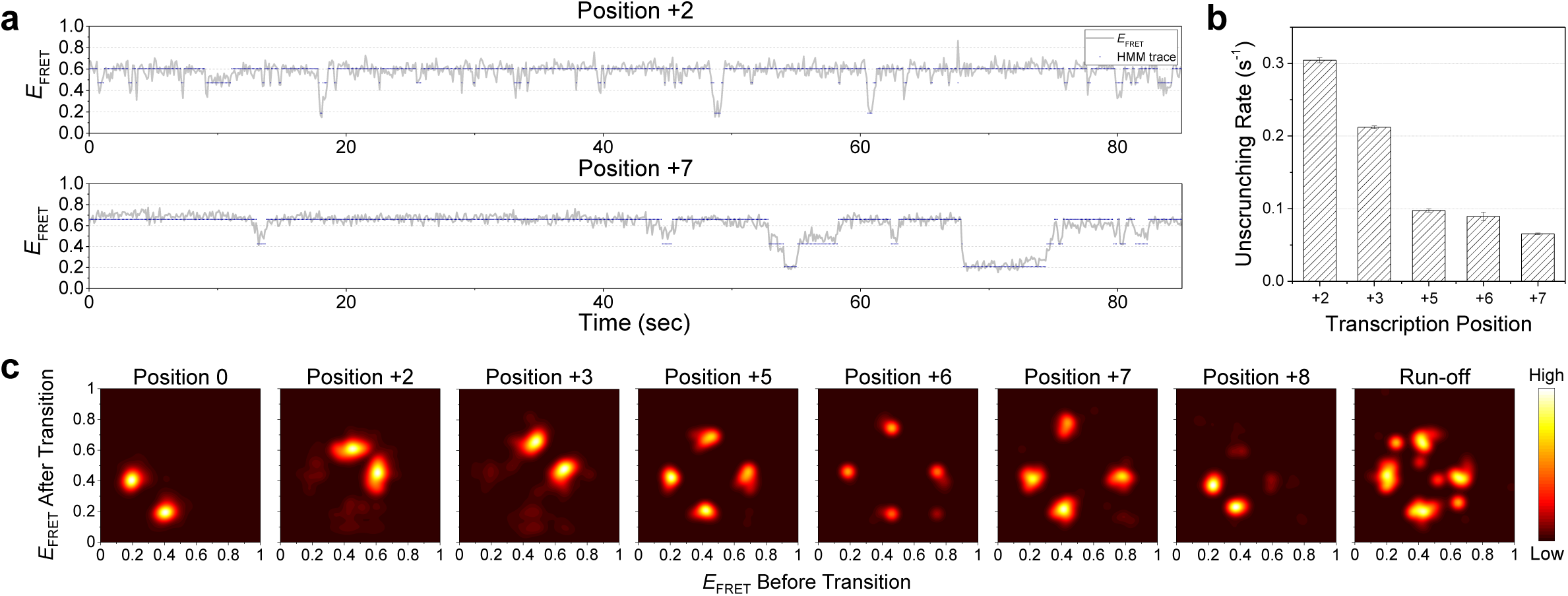
Hidden Markov analysis of smFRET traces throughout the initiation and elongation stages. **a**, Representative smFRET traces (gray) at positions +2 and +7 shown alongside hidden state traces (navy) from hidden Markov modeling assuming three hidden states. **b**, Unscrunching rates obtained from the hidden Markov analysis at each stalling position during initiation. **c**, Transition density plots from hidden Markov analyses of traces at each stalling position. 509, 184, 162, 45, 98, 51, 67, and 27 traces were used for positions 0, +2, +3, +5, +6, +7, +8, and run-off conditions, respectively.

To quantitatively assess the non-equilibrium transcription dynamics, we constructed FRET population maps by synchronizing smFRET traces at the moment when the FRET level exceeded 0.5 for the first time (time zero) (Fig. 2b). The FRET population map from flow-in measurements for progression to position +7 clearly showed that the low and mid FRET states coexisted before time zero, and the high FRET state dominated afterwards. Notably, the mid FRET population was dominant over the low FRET population immediately before time zero, indicating that the open promoter complex, but not the closed one, can proceed to transcribe and scrunch the DNA. After crossing 0.5, the high FRET state persisted for up to 10 seconds, reflecting the conformational stability of scrunched IC7. By contrast, the FRET population map from flow-in for progression to position +8 revealed an earlier transition from the high FRET state back to the mid or low FRET state, as demonstrated by the coexistence of these states after time zero (Fig. 2b). The FRET population map from flow-in for run-off revealed a similar behavior to that seen in the 0 to +8 measurements, but with the low FRET state dominating more at the end, possibly due to differences in the efficiency of incorporating 3’dCTP and CTP.

The dominant FRET population at position +8 and under run-off conditions was indistinguishable from that of bare DNA (Fig. 1c and 2b), possibly because DNA templates I/II happened to have similar *R*_D-A_ values for bare DNA and the elongation complex. This finding made it difficult to judge if the observed high-to-low FRET transition represented initiation-to-elongation transition or abortive initiation. To answer this question, we designed another DNA template +11/−11 that had the same sequence as DNA template II but different labeling positions, i.e., at positions −11 of the non-template strand and +11 of the template strand to enable better resolution in the low FRET range (Fig. 2c). Using this template, bare DNA and the open promoter (IC0) showed FRET levels of 0.32 and 0.75, respectively (Fig. 2d). IC7 showed a major FRET level of 0.64, which was lower than that of the open promoter; this finding is reasonable if we consider twisting of the downstream DNA arm, which would position the dye away from the transcription site at this stalling position. In addition, IC7 showed frequent transitions to a low FRET state, whose level matches that of the closed state in IC0 (Fig. 2d, Supplementary Fig. 3). The TIC stalled at position +8 showed a dominant FRET level of 0.44, which was different from that of bare DNA, and stayed at this level for a long time, just as the TIC at position +8 on DNA template II displayed a prolonged low FRET state (Fig. 2d, Supplementary Fig. 3). These results confirm that the low FRET state observed at position +8 with DNA template II is distinct from the low FRET state of bare DNA. Thus, we conclude that the TIC switches to the elongation complex at position +8 (EC8), with a large and abrupt change of conformation. As expected, under run-off conditions, DNA template +11/−11 displayed recovery of the FRET level to that of bare DNA, with a small remaining FRET population representing the TIC at intermediate steps (Fig. 2d).

### The transcription initiation bubble collapses upon transition to elongation

To obtain further support for our proposal that the transcription complex at +8 represents the elongation complex, we measured the kinetics of initiation bubble collapse. Figure 3a shows a model in which the initiation region from −4 to −1 remains melted up to position +7, and the initiation bubble collapses upon transition to elongation at position +8, reannealing the −4 to −1 region into duplex DNA (Fig. 3a). To monitor the transition from initiation to elongation, we substituted an adenosine at position −4 of the non-template strand with a 2AP residue, which emits fluorescence at 370 nm that is quenched when it pairs with a thymine residue (see online Methods)^36^. The fluorescence intensity of 2AP at position −4 was high in the open complex as the −4 basepair was melted, which was expected to decrease upon collapse of the initiation bubble. Thus, real-time monitoring of the fluorescence intensity during an *in vitro* transcription assay on two walking templates (Fig. 3b) allowed us to measure how the kinetics of initiation bubble collapse depends on the transcription position.

Upon walking to position +7, the fluorescence of 2AP remained constant for the total observation time (Fig. 3c). However, upon walking to position +8, the fluorescence of 2AP decreased to a lower level at a rate of 0.004 s^−1^. Similarly, upon walking to positions +9 and +10, the fluorescence intensity decreased to even lower levels at higher rates of 0.0165 s^−1^ and 0.027 s^−1^, respectively (Fig. 3c and 3d). These results are consistent with our interpretation of the single-molecule data that the complex at position +7 is an initiation complex with an open upstream bubble, and complexes from position +8 onward represent elongation complexes. The results also imply that the rate of transition to elongation increases with increasing length of the RNA/DNA hybrid from +8 to +10.

The progression of the major TIC conformation observed here is consistent with the known crystal structures of T7 and the human mitochondrial transcription machinery (Fig. 3e). The crystal structure of the yeast mitochondrial transcription machinery has not been determined; therefore, to represent IC0, we modeled the TIC structure of human mitochondrial RNAP (POLRMT) in complex with transcription factors (TFAM and TFB2M) (PDB: 6ERP)^41^. The promoter DNA in the TIC shows a sharply bent conformation, leading to a decrease in distance between the FRET dye pair at positions +16 and −16. For IC7, we modeled the known structure of the bacteriophage T7 RNAP in complex with the promoter DNA (PDB: 3E2E)^37^, which exhibits a further bent conformation, consistent with our smFRET data. For the elongation complex, we modeled EC9 of the human mitochondrial transcription complex with DNA (PDB: 4BOC)^38^, which shows a relaxed conformation of DNA.

### Landscape of conformational dynamics during transcription initiation and elongation

Different FRET states representing unbent, bent, and scrunched TIC conformations were well distinguished by our FRET histograms and traces (Fig. 1c and 1d). We used hidden Markov modeling (HMM) to extract kinetic information from the smFRET data (see online Methods)^43^. Figure 4A shows representative smFRET traces of IC2 and IC7 along with hidden state dynamics from the HMM analysis, assuming three hidden states. The high FRET state (scrunched conformation) frequently switched to the mid FRET state (unscrunched conformation) or the low FRET state (closed promoter), but mainly transitioned through the mid FRET state to reach the low FRET state, consistent with what we observed in synchronized FRET population maps (Fig. 2b). It is also worth noting that the unscrunching events were rare in IC7 compared with IC2, reflecting higher stability of the scrunched TIC at a later stage of transcription. The unscrunching rate revealed by the HMM analysis was 0.31 at position +2, and gradually decreased to 0.065 at position +7 during the initiation phase (Fig. 4b).

Next, we constructed transition density plots (TDPs) from the HMM analysis at each stalling position (Fig. 4c). The TDP at position 0 clearly shows the dynamics of promoter opening and closing (transition between low and mid FRET states). In IC2, the dominant events are the scrunching-unscrunching events (transition between mid and high FRET states) with rare promoter opening-closing events. After proceeding to positions +3, +5, +6, and +7, the level of the high FRET population gradually increased, consistent with the corresponding FRET histograms. At positions +5, +6, and +7, the relative density of the low-mid FRET transition became higher, which was due to a reduction in mid-high FRET transition events at later stages. At position +8, the high FRET population disappeared almost completely and low-mid FRET transitions dominated. Under run-off conditions, diverse FRET states coexisted, and transitions between them resulted in a complex TDP. The transformations of the TDP throughout the initiation and elongation stages highlight the transforming landscape of conformational dynamics throughout the early stage of transcription.

### The stalled initiation complex makes unscrunching transitions without dissociating RNA transcript

A straightforward interpretation of the frequent drops in the FRET level for stalled initiation complexes at positions +2 to +7 is that they represent dissociation of the RNA transcript, followed by unscrunching of the DNA (high-to-mid FRET transition), i.e., abortive initiation, and then the re-initiation of transcription using fresh NTP molecules (mid-to-high FRET transition), occasionally intervened by full closing and re-opening of the promoter (transitions to low FRET level). As we constantly supplied NTP mix to keep the reaction at equilibrium in the experiments described above, it was not clear if the drops and rises in the FRET level truly represented the abortion and re-initiation of transcription. Thus, we performed a different set of experiments in which we washed out the NTP mix, Rpo41, and Mtf1, and tracked changes in the FRET distribution over time. At position +2, in just 1 min after NTP wash-out, the scrunched population (high FRET) disappeared almost completely, suggesting rapid dissociation of the dinucleotide transcript (Fig. 5a and 5b). By contrast, at position +7, the scrunched population decreased much slower, with a half-life of approximately 5 min, reflecting the higher stability of IC7 compared with IC2 (Fig. 5a and 5c). This finding is consistent with the fact that exit of RNAP from the late initiation stage is markedly slower than that from earlier stages during bacterial transcription^35^.

**Figure 5.**
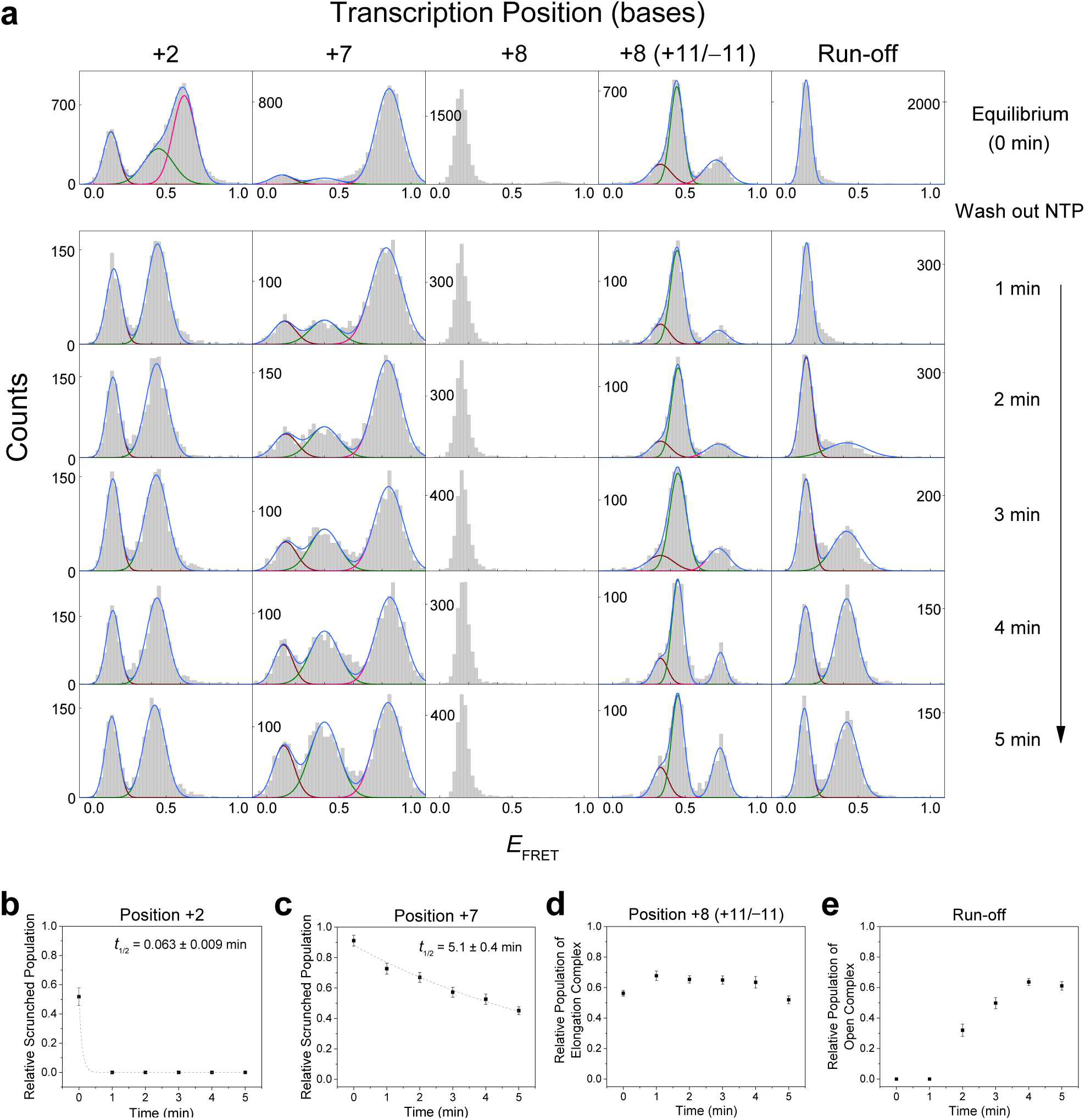
The rate of abortive initiation sharply depends on the transcription position. **a**, FRET histograms obtained at equilibrium and after washing out the NTP mixture for positions +2, +7, and +8, and run-off conditions. For position +8, results from DNA template +11/−11 are included to distinguish between the populations of the elongation complex and DNA only. Each histogram was obtained from 12 short movies taken during each minute after washing out the NTP mixture. **b**,**c**, The relative population of the high FRET state (scrunched complex) obtained from histograms at positions +2 (**b**) and +7 (**c**) shown relative to time. Each graph was fit to a single exponential decay curve, and the half-life is shown. **d**, The relative population of the mid FRET state (elongation complex) obtained from histograms at position +8 on DNA template +11/−11 shown relative to time. **e**, The relative population of the mid FRET state (open complex) obtained from histograms under run-off conditions shown relative to time.

DNA template II could not distinguish EC8 from the bare DNA at position +8, so we used DNA template +11/−11 to examine conformational changes at this position. Upon NTP wash-out, the scrunched population at position +8 diminished even slower than it did at position +7, demonstrating the high stability of EC8, which is typical of the elongation complex (Fig. 5a and 5d)^44^. To examine conformational changes under run-off conditions, we traced the open promoter population in DNA template II (Fig. 5a and 5e). The closed conformation dominated at equilibrium, indicating that once the open TIC was formed, transcription occurred relatively fast compared to the process of protein assembly and promoter opening. After NTP wash-out, the open promoter population gradually increased and saturated at a ratio of 0.6, which reflects the equilibrium ratio of the open promoter population in IC0, and the time delay indicates the time required for abortive initiation at mixed positions plus the time for new protein binding.

Next, we examined whether the conformational dynamics of the TIC disappeared in the absence of NTP mix to re-initiate transcription with. The NTP mix was washed out from the TIC equilibrated at position +7 and, in most traces, the FRET level dropped a little while after the wash-out, possibly representing an abortive initiation event. However, to our surprise, many traces showing a drop in the FRET level subsequently returned to the high FRET level (Fig. 6a, Supplementary Fig. 5), which must have happened without dissociating RNA strands and synthesizing new ones because there was no NTP substrate remaining. This observation demonstrates that the TIC makes conformational transitions not necessarily by aborting RNA synthesis. As such dynamics occurred with bound RNA strands, the observed mid or low FRET level should not represent the conformation of the open promoter or the bare DNA template.

**Figure 6.**
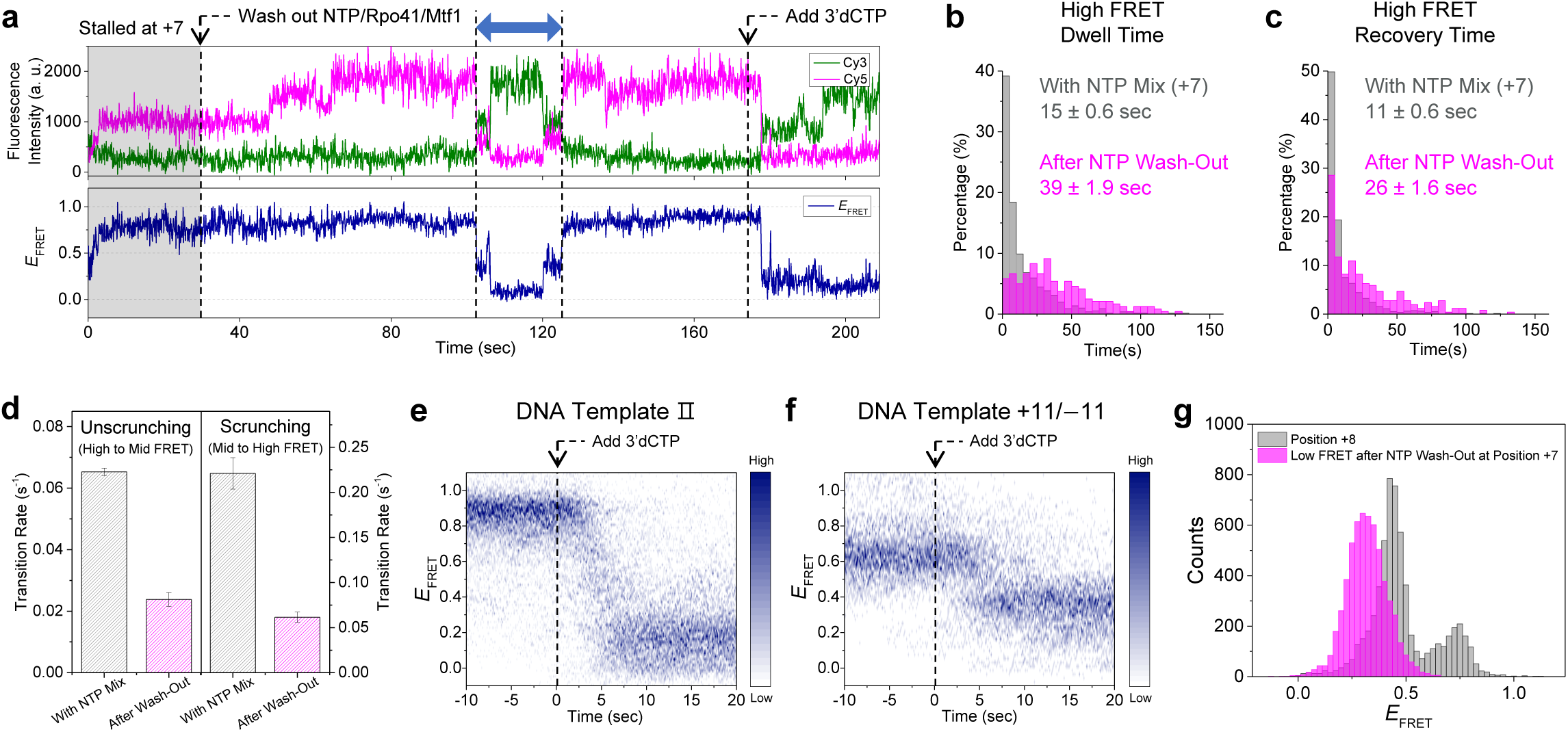
The stalled initiation complex makes conformational transitions without dissociating the RNA transcript. **a**, Representative smFRET trace showing the conformational dynamics of the TIC of DNA template II equilibrated at position +7 (gray region; 0.5 mM each of ATP, GTP, and UTP) after washing out the NTP mix (first arrow). Subsequently, 0.5 mM 3’dCTP was added to promote progression to position +8 (second arrow). The abrupt drop in the FRET efficiency indicates successful progression to position +8. **b**, Dwell time histogram of the high FRET (scrunched) state in the presence of NTP mix (grey; 809 events) and after NTP wash-out (magenta; 214 events). **c**, The time taken to recover the high FRET state after dropping to lower FRET levels (blue arrow in (**a**)) in the presence of NTP mix (grey; 843 events) and after NTP wash-out (magenta; 284 events). **d**, Comparison of unscrunching and scrunching rates in the presence of NTP mix and after NTP wash-out, measured as the inverse of average dwell times in (**b**) and (**c**). **e**,**f**, FRET evolution maps generated from the traces supplied with 0.5 mM 3’dCTP after long stalling at position +7 by washing out the NTP mix, synchronized at the moment of flowing in 3’dCTP (0 seconds). 53 and 41 traces were used for the maps generated for DNA templates II and +11/−11, respectively. **g**, Using DNA template +11/−11, FRET histogram of low-FRET events after NTP wash-out at position +7 (magenta) was compared to that of position +8 (grey, from Fig. 2d).

Compared with that in the presence of NTP, the dwell time of the high FRET state in the absence of NTP was markedly longer and displayed a non-exponential distribution, further supporting the proposal that the dynamics of conformational changes in the absence of NTP are distinct from those of abortive initiation (Fig. 6b). The duration that the TIC spent out of the high FRET state was also much longer in the absence of NTP than in the presence of 0.5 mM NTP (Fig. 6c). As the majority of NTP-independent transitions occurred to and from a mid FRET level (Supplementary Fig. 5), we measured the high-to-mid and mid-to-high transition rates as if they represented unscrunching and scrunching kinetics, respectively, occurring with bound RNA strands. In the absence of NTP, both transition rates were much lower than those in the presence of NTP, indicating that in the absence of NTP, the TIC goes through slower transitions that are distinct from those of abortive transcription and re-initiation (Fig. 6d).

After stalling the TIC at position +7 without NTP for longer than 2 min, we flowed in 0.5 mM 3’dCTP to see if the TIC could still progress to the elongation stage. Soon after the flow-in, the FRET level dropped to that of EC8 (Fig. 6a, Supplementary Fig. 5). This occurred consistently in most traces and a FRET population map generated from the traces synchronized at the moment of 3’dCTP flow-in clearly showed the high-to-low FRET transition occurring in 5 seconds (Fig. 6e). As DNA template II could not distinguish the conformation of EC8 from that of bare DNA, we repeated the same measurement using DNA template +11/−11, and the majority of the traces displayed transitions to the FRET level of EC8 (Fig. 6f). These results show that the TIC can be stalled at a late stage of initiation, make multi-step conformational transitions to unbent or unscrunched DNA conformations without completely dissociating RNA strands, and then resume transcription to progress to the elongation stage.

As shown above, the transient drop of FRET level in stalled IC7 should not represent unscrunching of IC7 by abortive dissociation of RNA. Another possibility is that it represents temporary transition forward to an EC-like structure where the upstream promoter is released prior to the addition of the 8^th^ nucleotide and the promoter is unbent. FRET histogram was built from low-FRET events in IC7 with DNA template +11/−11 after NTP was washed out. Then it was compared to the FRET histogram of EC8 with DNA template +11/−11 (Fig. 6g). Major FRET level of the transient drops was distinguishably lower than that of EC8, suggesting that it represents a structure distinct from EC8. Thus, we conclude that IC7 branches into a conformation, which represents neither unscrunching by RNA dissociation nor an EC-like structure. 30% of smFRET traces from stalled IC7 showed transient drops of FRET level while 64% showed an irreversible drop (Sup. Fig. 6). Branching ratio between them is in rough agreement with the ratio found from the decay rates of high FRET state before and after NTP wash-out (0.026 vs (0.067−0.026); Fig. 6b). The only possibility is that this distinct transient conformation represents an unscrunched downstream DNA that still has an RNA bound, which can be elongated further. Consequently, downstream DNA unscrunching would destabilize the RNA-DNA hybrid and fray the 3’-end of the RNA, resembling a backtracked complex.

## Discussion

In this study, we combined single-molecule techniques with ensemble biochemical assays to dissect the conformational dynamics of the mitochondrial TIC throughout the initiation and elongation stages. At each nucleotide step, the TIC did not adopt a stationary conformation, but rather exhibited dynamic transitions between closed, open, and scrunched conformations. These dynamics persisted but gradually diminished along the initiation stage, culminating in stabilization of the scrunched conformation toward the end of the initiation stage. The downward FRET transitions of the scrunched TIC represent not only abortive initiation events, accompanied by the dissociation and new synthesis of RNA strands, but also pure conformational dynamics without the dissociation of RNA strands. The rate of abortive initiation at position +7, measured as the rate at which the scrunched population decreased after washing out NTP mix (0.0033 s^−1^; Fig. 5c) was much lower than the unscrunching rate determined via a hidden Markov analysis (0.065 s^−1^; Fig. 4b), indicating that a large part of the downward FRET transitions were not accompanied by RNA dissociation.

This finding is reminiscent of the NTP-independent scrunching-unscrunching dynamics of bacterial transcription systems^3^. The unscrunching motion in bacterial transcription mechanistically resembles backtracking that leads to long-lived catalytically inactive initiation complexes; this process is thought to play a regulatory role by maintaining an “elongation-ready” initiation complex that can be conditionally triggered. Backtracking during transcription is known to occur by extruding the 3’ end of nascent RNA through the secondary channel^45^. There is yet to be any evidence that mitochondrial RNAP has a secondary channel like multi-subunit RNAPs. The absence of a distinct secondary channel in mitochondrial RNAP might explain why the lifespan of the unscrunched state at position +7 in the Rpo41-Mtf1 complex was relatively short compared to what was found in the bacterial system. However, the structure of human mitochondrial RNAP suggests that there is room for 3’-end extrusion near the active site. Thus, the NTP-independent unscrunching motion may represent transient backtracking during initiation that could regulate transcription efficiency. Future experiments on the basepairing state of RNA and the unscrunched structure of TIC would reveal the identity of the transiently unscrunched state found here.

Hidden Markov analysis of single-molecule traces allowed us to quantify the conformational transition rates. The unscrunching rate decreased more than 4-fold as transcription initiation proceeded from position +2 to +7, implying that the conformational stability of scrunched DNA is highly sensitive to changes in the TIC structure. The sharp dependence of the unscrunching rate on the transcript length might stem from differences in the stability of RNA-DNA hybrid, which may provide a mechanism to prevent incorrectly transcribed RNA primers entering the elongation stage due to lower stability of the RNA-DNA hybrid. It further relates to the observation that mitochondrial transcription efficiency varies up to ∼ 100-fold between different initiating nucleotide sequences at positions +1 and +2^10^, suggesting that the scrunched TIC conformation is differentially stabilized by different initiating sequences. Taken together, these results suggest that the initiation stage may function as a regulatory step to control transcription kinetics depending on promoter sequences and biochemical circumstances. In bacteriophage and bacterial transcription systems, the progression through transcription initiation serves as a rate-determining step in transcription^3,30^.

It is intriguing that the conformational progression of the TIC within the initiation stage was more reversible in this two-protein transcription system of yeast mitochondria than in the more primitive, single-protein transcription system of T7 bacteriophage. While transcription initiation by T7 RNAP shows a single dominant FRET population at each step representing one stable conformation^30^, Rpo41 and Mtf1 showed highly reversible dynamics between distinct conformations. Rpo41 is homologous to T7 RNAP, but requires Mtf1 to initiate transcription^13,14,46^. Mtf1 is thought to stabilize the open promoter complex^18,24,36^, but our results suggest that Mtf1 in complex with Rpo41 allows dynamic exchange between scrunched and unscrunched conformations, which might result in enhanced production of abortive transcripts. This proposal resonates with the observation that the addition of Mtf1 increases the production of abortive transcripts by Rpo41 on pre-melted DNA^18^. Thus, our observations suggest novel roles of initiation factors in facilitating conformational transitions of the TIC, thereby providing proofreading steps or conditional gates that determine progression to the elongation stage.

Transition to the elongation stage was found from smFRET measurements to occur in a single step precisely at position +8, as indicated by a large change in the FRET level and the disappearance of conformational fluctuations, setting the TIC in a stable and less bent conformation. This contrasts with the smFRET observation on the single-subunit T7 transcription system, in which the transition occurs gradually over positions +8 to +12^30,39^. Ensemble-level biochemical studies and crystal structure analyses have also suggested that transition of T7 RNAP to the elongation stage occurs in multiple steps with a series of conformational transitions, producing abortive transcripts of at least up to 20 bases^37,39,40,47^. The interplay between RNAP and initiation factors in yeast mitochondria appears to make an abrupt release of the promoter at a sharply defined position, which is not reversible. This process prevents the elongation complex from reversing to the initiation stage or escaping through an abortive pathway. Irreversible dissociation of Mtf1 in the elongation stage possibly functions as a kinetic latch to prevent such non-productive pathways. Though Mtf1 is known to dissociate after transcribing 13 nucleotides^48^, it is not clear whether it dissociates at or after the unbending transition at position +8. On the other hand, 2AP measurements showed the collapse of the initiation bubble starting at position +8, but the bubble appeared to gradually collapse over positions +8 to +10 (Fig. 3c). In case of T7 RNAP, promoter release and bubble collapse occur synchronously. This does not appear to be the case in mitochondrial transcription, where the TIC takes further steps until the initiation bubble fully zips up. Our results suggest that promoter release occurs first to form an unbent EC-like state and then bubble collapse follows to complete transition to elongation.

Integrating the conformational transitions identified in this study, we constructed a kinetic model of mitochondrial transcription initiation (Fig. 7). Initial bound complex of DNA/Rpo41/Mtf1 transitions between closed and open promoter states (IC0_closed_ and IC0_open_). Upon incorporation of the initiating nucleotides, IC0_open_ starts scrunching in stepwise manner (ICn_scrunched_), but it often reverts to IC0_open_ and even to IC0_closed_ by dissociating RNA to re-initiate transcription. The rate of abortive transcription greatly decreases with increasing length of RNA, making the scrunched TIC increasingly stable. In addition to the abortive transcription, IC7_scrunched_ makes slower, reversible transitions to an unscrunched conformation (IC7_unscrunched_), which may also occur at earlier positions. In these states, the transcript remains bound to the complex, possibly with its 3’-end extruding from the complex. Then, IC7_scrunched_ makes a large irreversible conformational transition at position +8 by releasing the upstream DNA that might be accompanied by dissociation of Mtf1 (EC8). It is followed by gradual zipping of the upstream bubble in the following steps.

**Figure 7.**
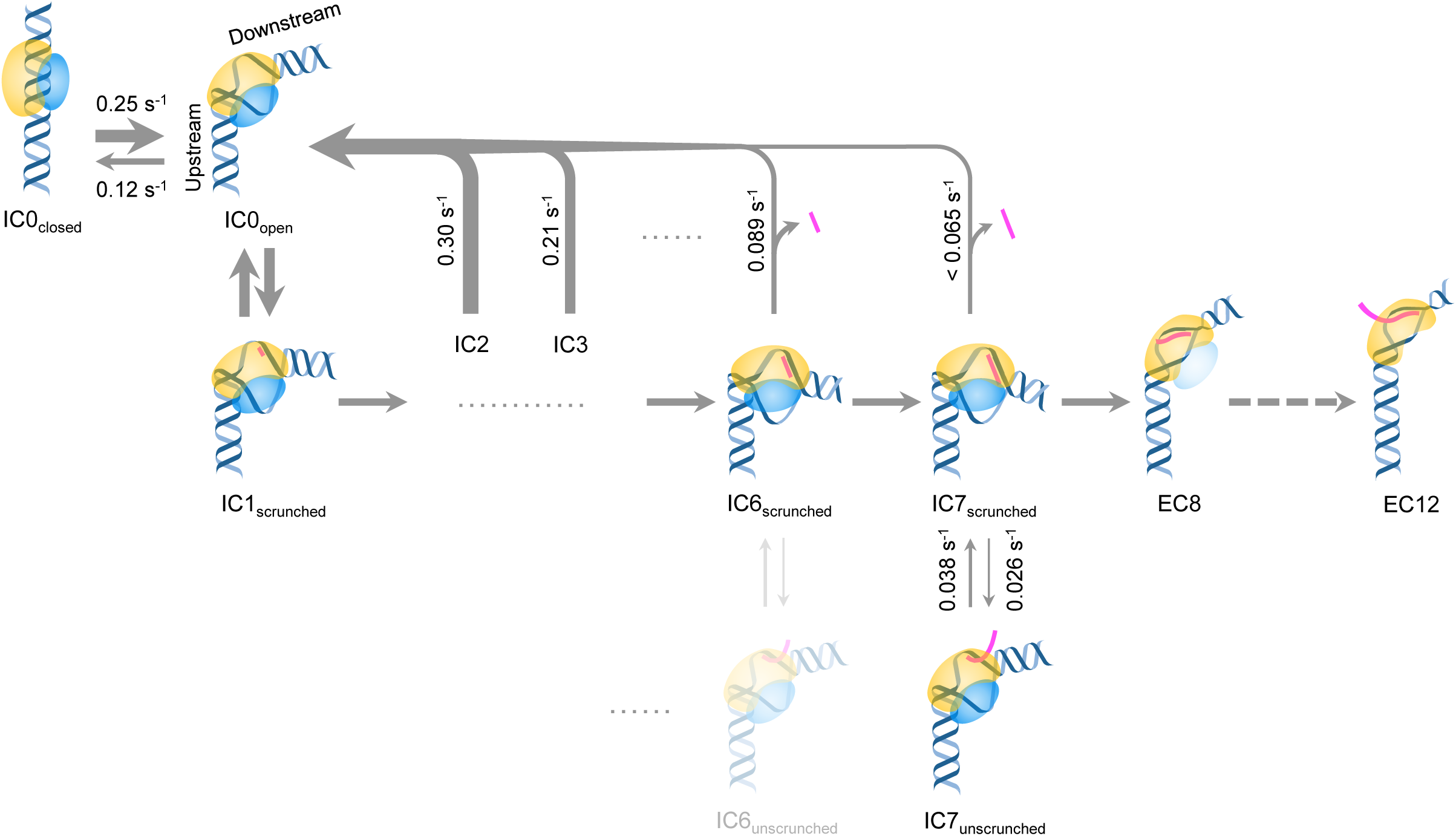
Kinetic model of mitochondrial transcription initiation. Conformational states and their transition kinetics identified in this study were integrated into the model. Measured transition rates were marked which were reflected in the thickness of arrows. Blank arrows represent indistinguishable transitions or transitions whose rates depend on NTP concentration. Unidentified states (IC6_unscrunched_) or molecules (Mtf1 in EC8) were expressed semi-transparent. DNA/RNA oligos and proteins were drawn as in Fig. 1a.

In conclusion, our study reveals the highly dynamic and reversible nature of mitochondrial transcription initiation. We propose that the dynamic initiation stage and irreversible transition to the elongation stage serve as key features that distinguish multi-subunit transcription machineries from simpler single-subunit systems. Our findings provide novel mechanistic perspectives on how transcription machinery regulates the progression of transcription initiation, and form a basis for studies of more sophisticated transcription machineries of higher organisms.

## Online Methods

### Preparation of DNA templates

DNA oligonucleotides were custom synthesized with biotin and amino modifications, and were purified by HPLC (Integrated DNA Technologies, USA). The oligonucleotides were fluorescently labeled at the amine groups by standard assays using Cy3 or Cy5 NHS esters (Lumiprobe, USA), and unreacted dyes were removed by ethanol precipitation. To generate duplex DNA molecules, single-stranded DNAs were mixed in a 1:1 ratio, annealed at 95°C for 1 min, and then slowly cooled to room temperature for 1 h.

### Single-molecule measurements

Single-molecule fluorescence signals were detected using a custom-built total internal reflection fluorescence (TIRF) microscope^49^. The sample surface was prepared by coating quartz slides (Finkenbeiner, USA) with a 40:1 mixture of mPEG-SVA (MW 5,000) and biotin-PEG-SVA (MW 5,000) (Laysan Bio, USA) after treatment with (3-aminopropyl)trimethoxysilane (Sigma, USA). The surface was coated with NeutrAvidin (ThermoFisher Scientific, USA) and DNA templates were immobilized via the biotin-NeutrAvidin interaction. Rpo41 (100 nM) and Mtf1 (100 nM) were flowed into the chamber and incubated with the DNA templates for 3 min. Excess unbound proteins were then washed away and imaging buffer (100 mM Tris-acetate pH 7.5, 50 mM potassium glutamate, 10 mM magnesium acetate, 0.6% glucose, 1 mg/ml glucose oxidase (from Aspergillus niger VII; Sigma, USA), 0.04 mg/ml catalase (from bovine liver; Sigma), ∼3 mM Trolox, and controlled concentrations of various combinations of NTPs) was added to stall the TIC at varying positions. Fluorescence movies of the donor and acceptor channels were recorded using an EMCCD camera (iXon Ultra 897; Oxford Instruments, UK). All measurements were performed at 25°C.

### Single-molecule data analysis

Movies obtained using the TIRF microscope were analyzed using custom software to extract single-molecule fluorescence traces, as described previously^6^. The FRET efficiency was calculated with background and leakage correction as *E*_FRET_ = (*I*_*A*_ − 0.08 × *I*_*D*_)/(*I*_*D*_ + *I*_*A*_), where *I*_*D*_ and *I*_*A*_ are the background-subtracted intensities of the donor and acceptor dyes, respectively. The acceptor dyes were briefly excited at the beginning and end of each movie to exclude traces lacking acceptor dyes from further analysis. Each FRET histogram was built from more than 50 movies by selecting traces with a single pair of Cy3 and Cy5 dyes and representing each trace by *E*_FRET_ averaged over five frames. Hidden Markov analysis was performed using ebFRET software developed by the Gonzalez group^43^.

### 2-Aminopurine fluorescence assay

Steady-state fluorescence measurements were carried out at 25°C using a Fluoro-Max-2 spectrofluorometer (Jobin Yvon Spex Instruments S.A., Inc., USA) in buffer containing 50 mM Tris-acetate (pH 7.5), 100 mM potassium glutamate, and 10 mM magnesium acetate. The fluorescence spectra of 200 nM 2AP-incorporated duplex promoters were collected from 350 to 420 nm (6 nm bandwidth) with excitation at 315 nm (2 nm bandwidth) after the sequential addition of Rpo41 (400 nM), Mtf1 (400 nM), and initiating +1 +2 NTPs (1 mM). After subtracting the contributions of the buffer and proteins in the presence of unmodified DNA, the corrected 2AP fluorescence intensities between 360 nm and 380 nm were integrated for comparison.

## Acknowledgements

This work was supported by the National Research Foundation of Korea (2017R1D1A1B03036239, 2017M3A9E2062181, and 2018R1A5A1024340) and the Institute for Basic Science (IBS-R022-D1) to H.K.; the National Institute of Health grant R35 GM118086 to S.S.P.; American Heart Association 16PRE30400001 and Louis Bevier Dissertation Completion Fellowship from Rutgers University to U.B.

## Author contributions

H.K. and S.P. conceived the project. B.S., U.B., S.L., H.C., J.S., and A.D. performed the experiments and analyzed the data. B.S., H.K., U.B., and S.P. wrote the manuscript.

