## Supplemental Figures for "The Dynamic Landscape of Transcription Initiation in Yeast Mitochondria"

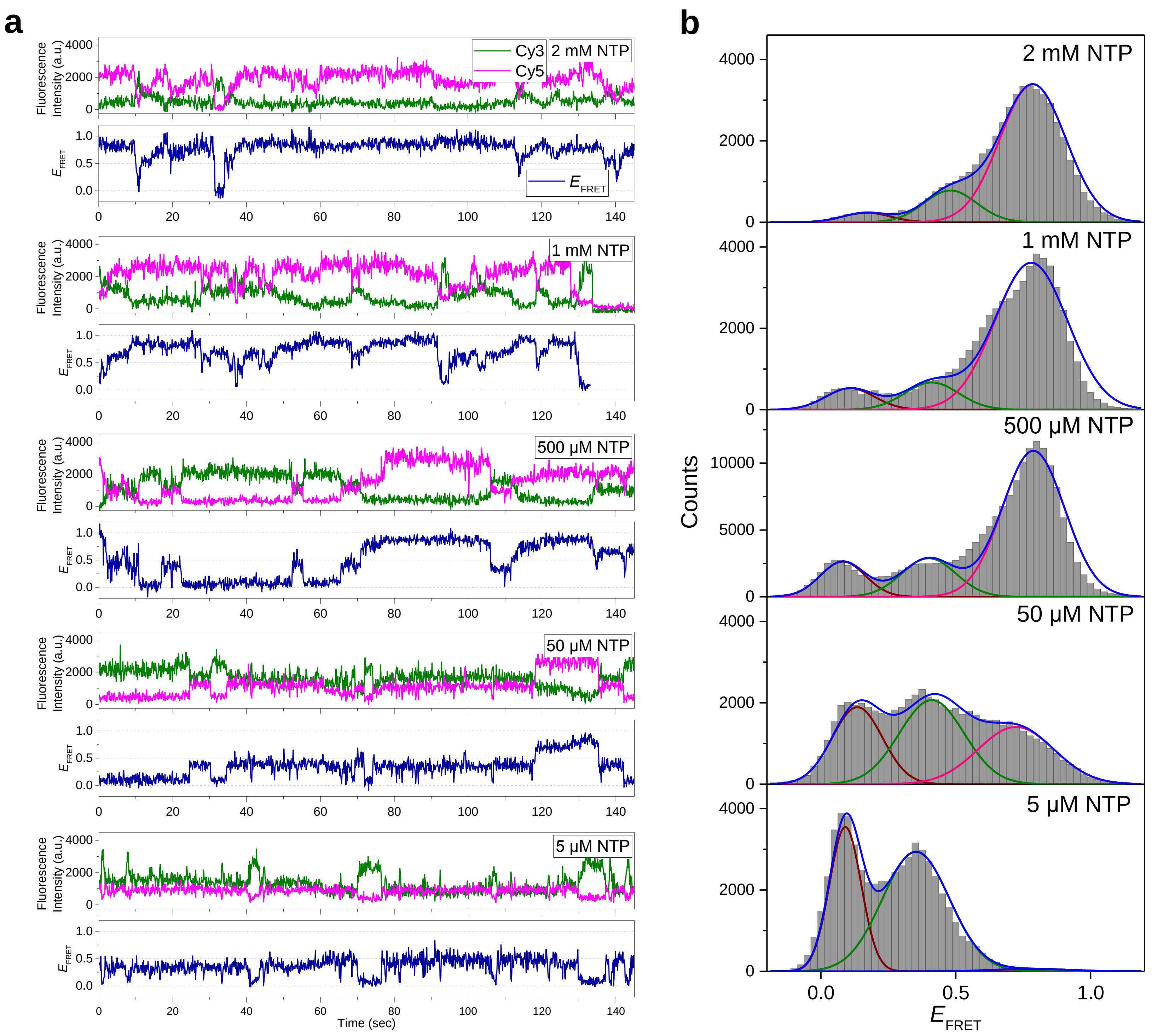


**Supplementary Figure 1. a**, Representative smFRET traces of the TIC at varying concentrations of NTP mix. The concentrations represent those for each kind of nucleotide. **b**, FRET histograms obtained at each NTP concentration. 45, 51, 153, 46, and 45 traces were used for the histograms at 2 mM, 1 mM, 500 μM, 50 μM, and 5 μM NTP, respectively. Each histogram was fit to two or three Gaussian peaks as in Figure 1c.

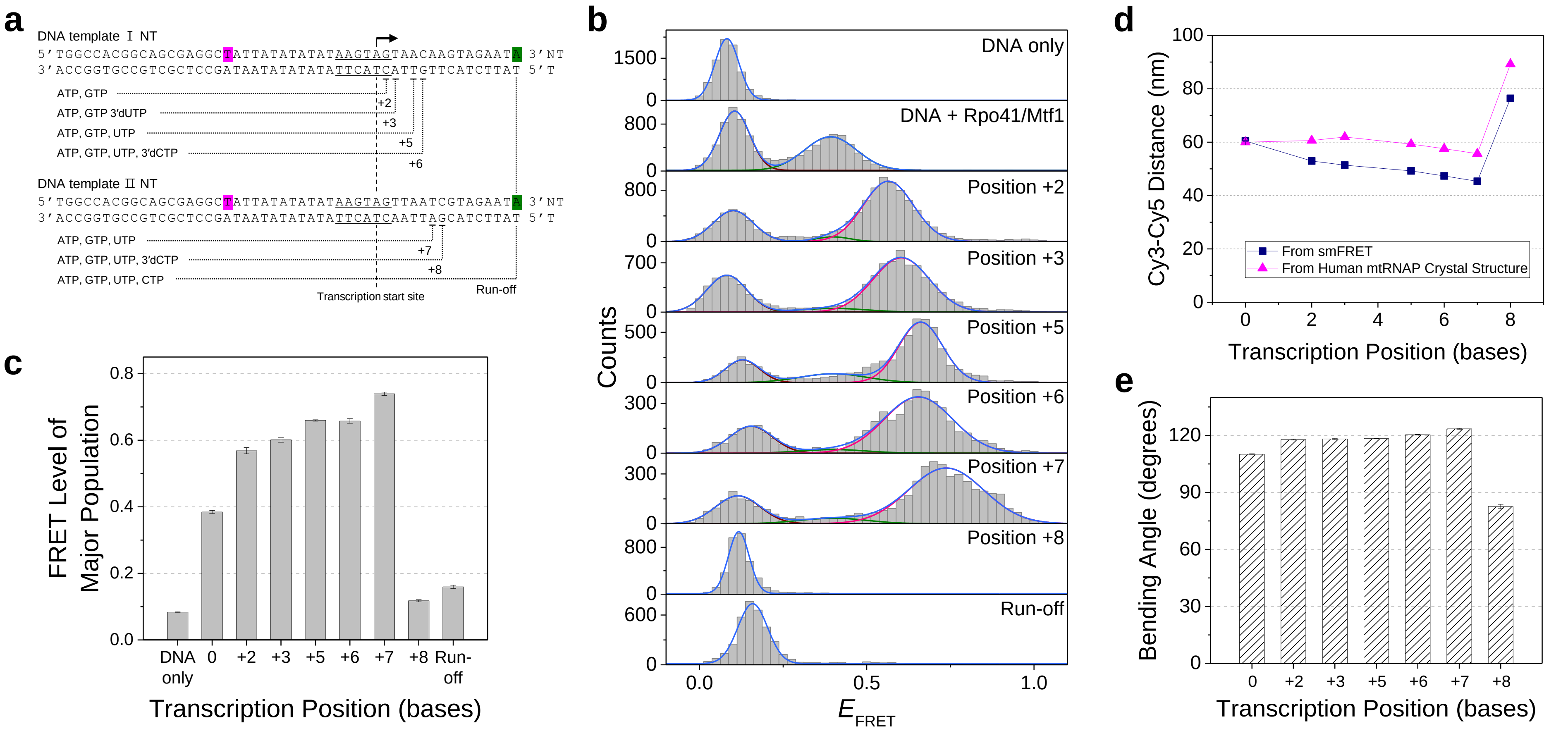


**Supplementary Figure 2. a**, Schematic design of the DNA templates used to measure the bending angles. The design is the same as that shown in Figure 1, with the exception that both Cy3 and Cy5 were located on the non-template strand at positions +16 and −16. **b**, FRET histograms and Gaussian fitting as described in Figure 1c. **c**, The FRET level of the major population in (**b**) shown for each stalling position as the center of the major Gaussian peak. **d**, Comparisons between the Cy3-Cy5 distances calculated from smFRET measurements and from the crystal structure of the human mitochondrial TIC (PDB: 6ERP). **e**, The bending angle of the DNA template calculated from smFRET data at each position. The error bars represent propagation of the errors in FRET levels.

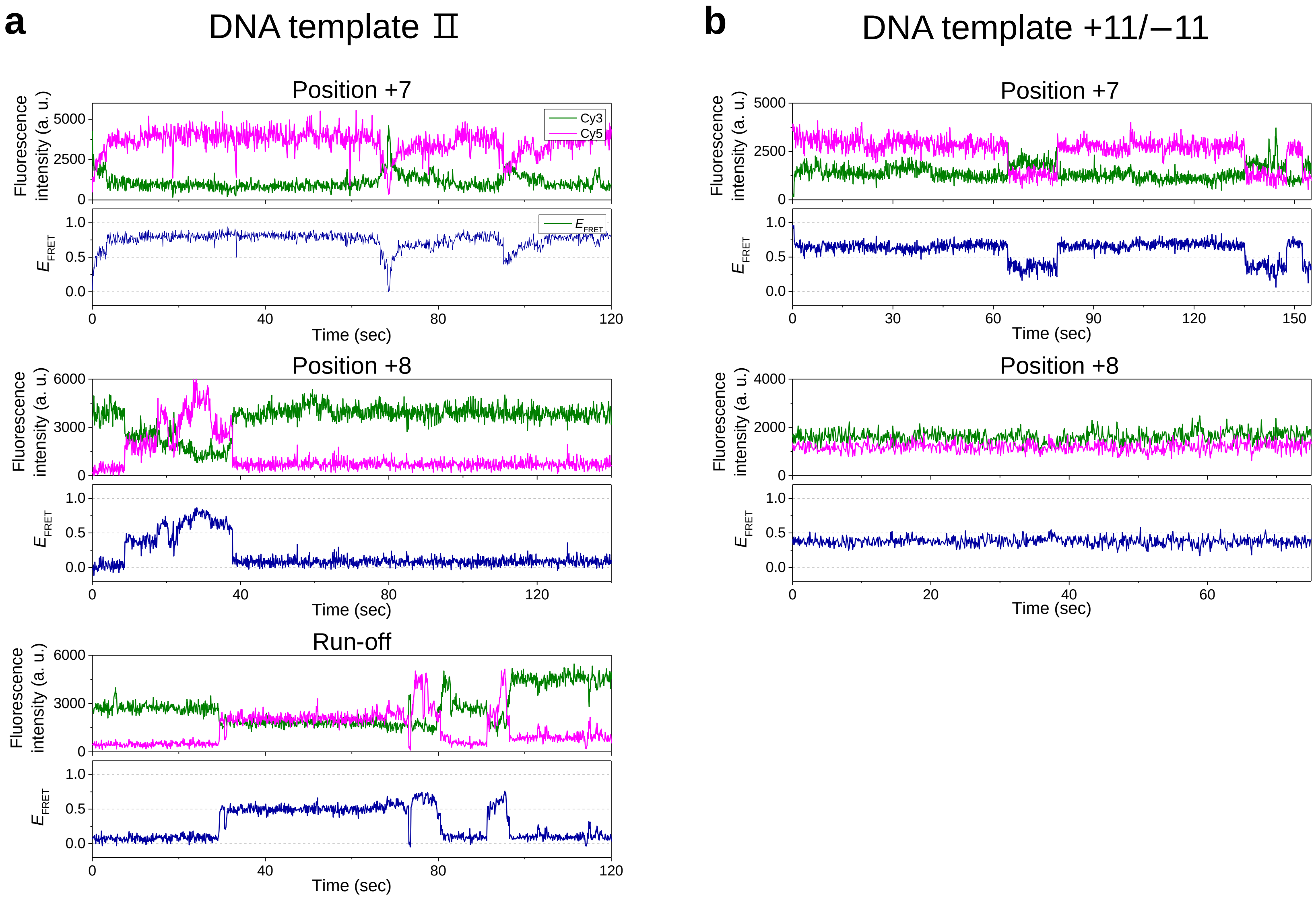


**Supplementary Figure 3. a**, Representative smFRET traces at positions +7 and +8 and under run-off conditions on DNA template Ⅱ. **b**, Representative smFRET traces at positions +7 and +8 on DNA template +11/−11.

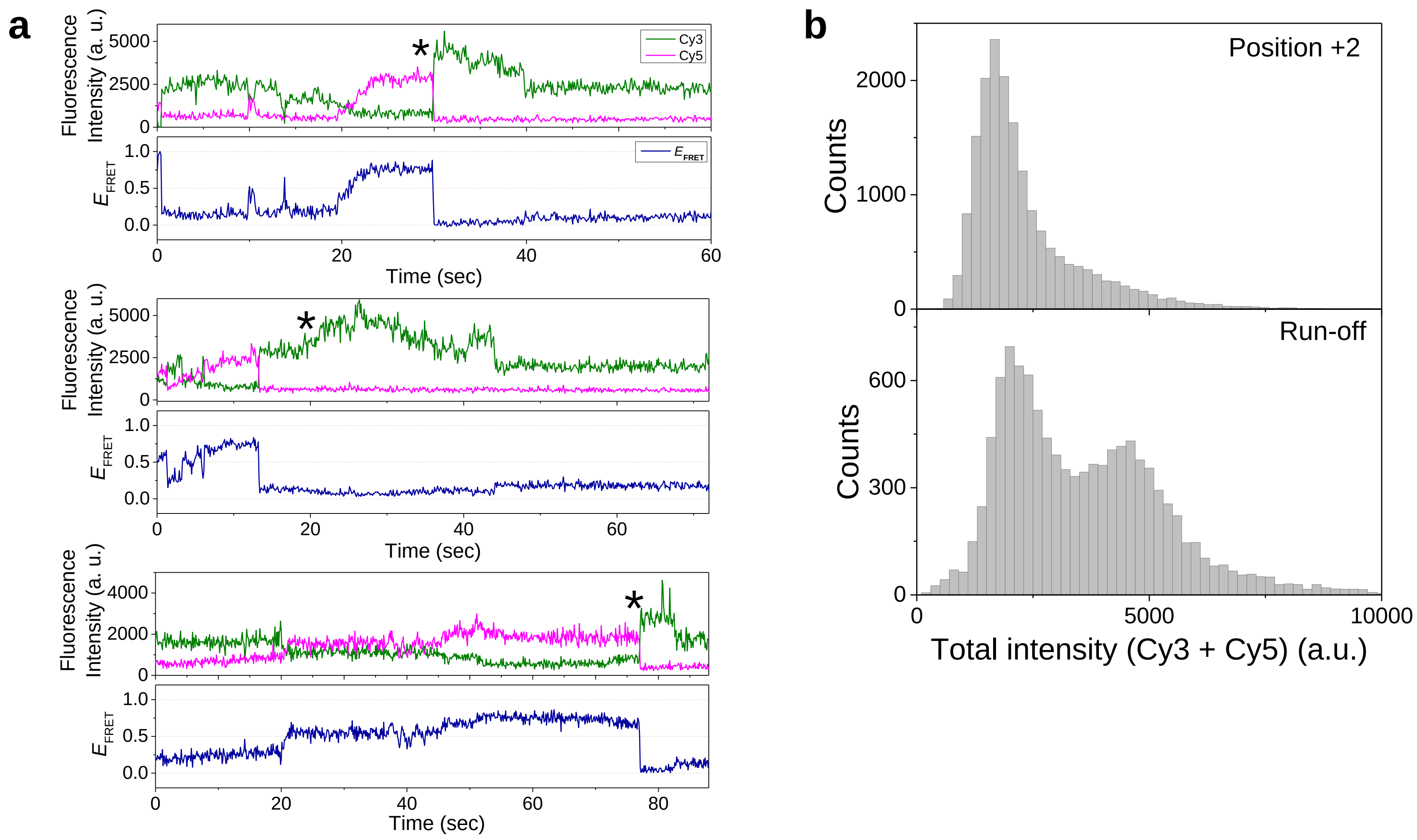


**Supplementary Figure 4. a**, Representative smFRET traces under run-off conditions for DNA template II, showing enhanced fluorescence signals (asterisks) following the drop in the FRET level. **b**, Histograms of total fluorescence intensities comparing position +2 and run-off conditions.

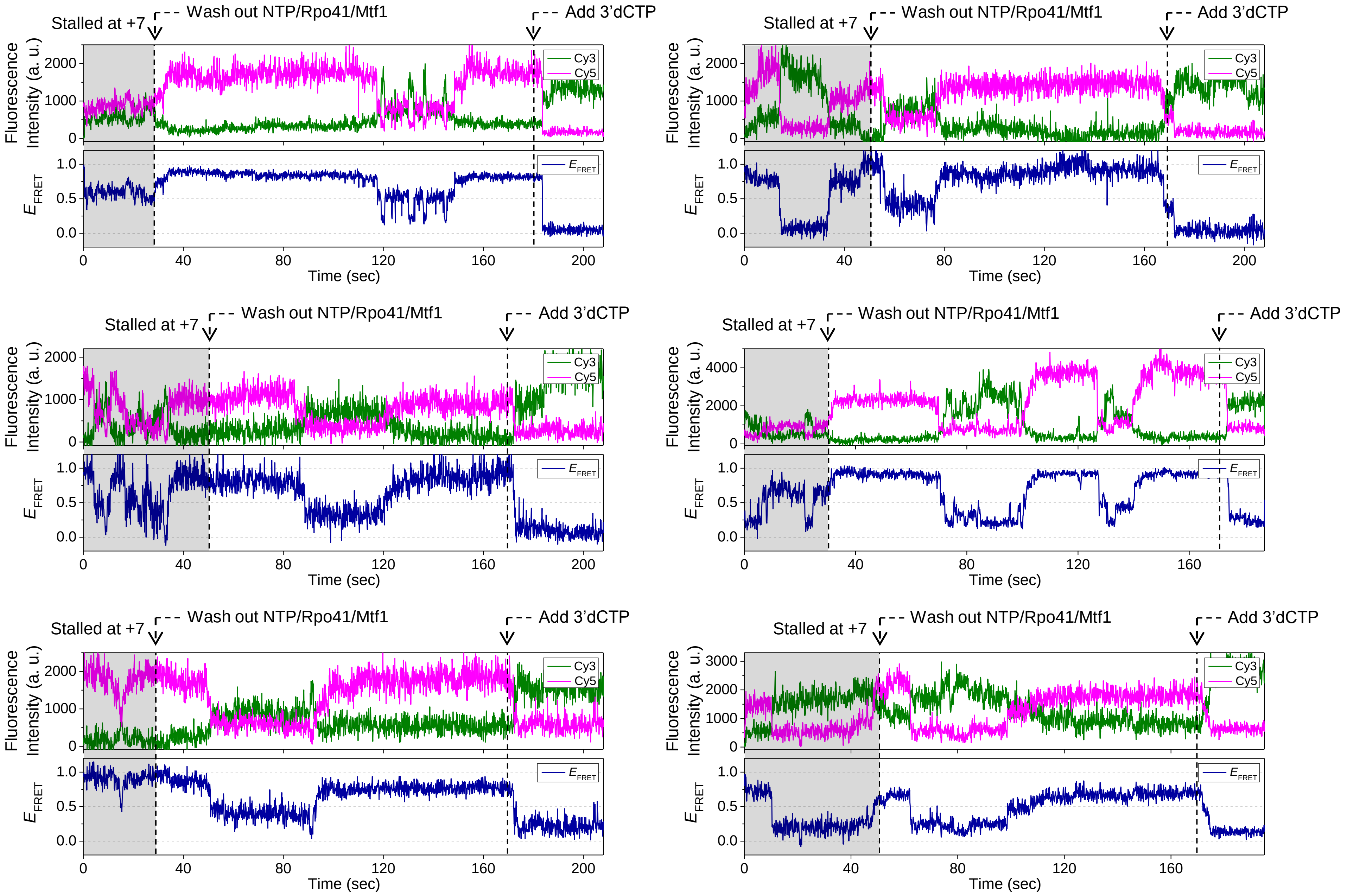


**Supplementary Figure 5.** Representative smFRET traces showing the conformational dynamics after washing out the NTP mix (first arrow) from the TIC of DNA template II equilibrated at position +7 (gray region; 0.5 mM each of ATP, GTP, and UTP). Subsequently, 0.5 mM 3'dCTP was added to promote progression to position +8 (second arrow). The abrupt drop in the FRET efficiency indicates successful progression to position +8.

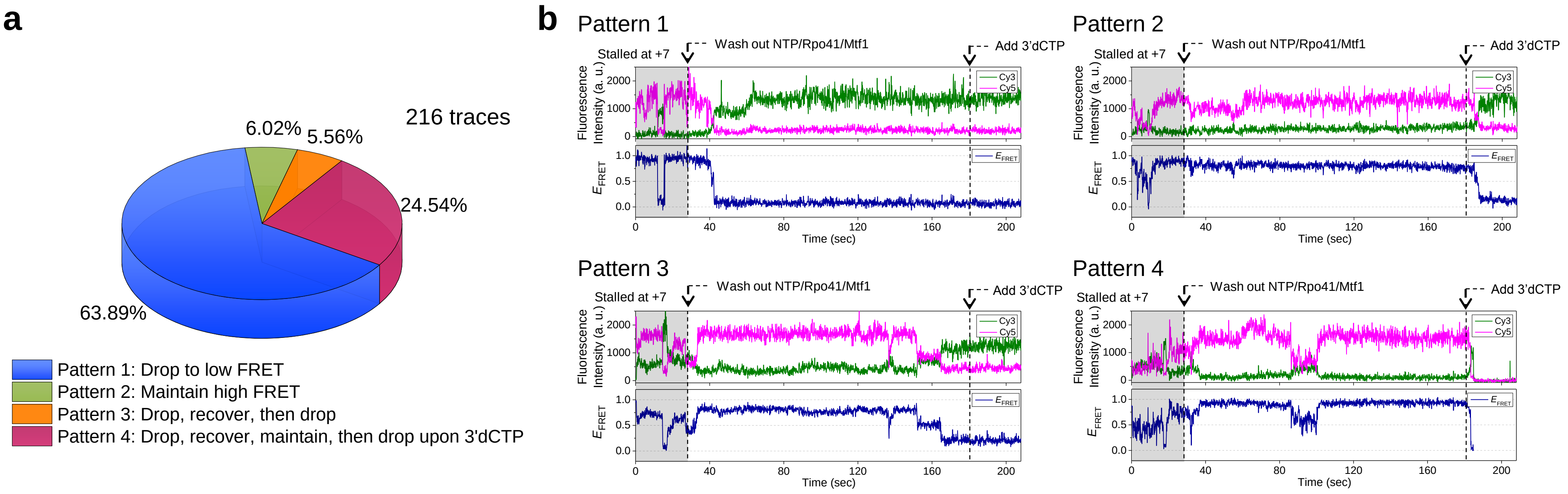


**Supplementary Figure 6. a**, Diagram of four distinct patterns of single-molecule traces observed in NTP wash-out experiments. Pattern 1: FRET level dropped and remained low after NTP wash-out. Pattern 2: FRET level remained at high FRET until flowing in 3’dCTP. Pattern 3: FRET level dropped and recovered back to high level, but dropped again before flowing-in 3’dCTP, preventing to check the capability of progressing to elongation. Pattern 4: FRET level dropped, recovered, and remained high until flowing-in 3’dCTP, showing the capability of progressing to elongation. Patterns 3 and 4 represent smFRET traces showing FRET level drop followed by the recovery of high FRET level. **b**, Example traces for each pattern of single-molecule trace.
